# The autism spectrum disorder risk gene *NEXMIF* alters hippocampal CA1 cellular and network dynamics

**DOI:** 10.1101/2022.10.21.513282

**Authors:** Rebecca A. Mount, Mohamed Athif, Margaret O’Connor, Amith Saligrama, Hua-an Tseng, Sudiksha Sridhar, Chengqian Zhou, Heng-Ye Man, Xue Han

## Abstract

Perturbations in autism spectrum disorder (ASD) risk genes disrupt neural circuit dynamics and ultimately lead to behavioral abnormalities. To understand how ASD-implicated genes influence network computation during behavior, we performed *in vivo* calcium imaging from hundreds of individual hippocampal CA1 neurons simultaneously in freely locomoting mice with total knockout of *NEXMIF. NEXMIF* is an ASD risk gene most highly expressed in the hippocampus, and *NEXMIF* knockout in mice creates a range of behavioral deficits, including impaired hippocampal-dependent memory. We found that *NEXMIF* knockout does not alter the overall excitability of individual neurons but exaggerates movement-mediated neuronal responses. At the network level, *NEXMIF* knockout creates over-synchronization of the CA1 circuit, quantified by pairwise correlation and network closeness centrality. These neuronal effects observed upon *NEXMIF* knockout highlight the network consequences of perturbations in ASD-implicated genes, which have broad implications for cognitive performance and other ASD-related behavioral disruptions.

## Introduction

Autism spectrum disorder (ASD) is a neurodevelopmental disorder that affects 1 in 54 children (by the age of 8) in the United States^1^. ASD is characterized by two core behavioral symptoms. The first is difficulty with social interaction and communication, and the second is repetitive, restrictive behaviors and interests^2^. ASD is often co-morbid with other medical conditions such as epilepsy, gastro-intestinal issues, sleep problems, and immune dysfunction^3^. As one of the most heritable neuropsychiatric disorders, the genetic bases of ASD are widely heterogenous and often polygenic^4^. Thus far, human genomic studies have identified numerous genes implicated in ASD risk. To understand the contribution of these genes to ASD pathophysiology, many transgenic mice^5,6^ and non-human primates^7^ containing such gene disruptions have been developed to model aspects of behavioral, molecular, and cellular phenotypes seen in individuals with ASD.

Many ASD risk genes are thought to disrupt neural network excitability by increasing excitatory/inhibitory (E/I) balance within neural circuits^8^. While there is little direct experimental evidence of how increased E/I ratio alters network dynamics, computational modeling has revealed that E/I balance is critical for maintaining proper asynchrony within a network^9^ and increased E/I ratio elevates neural synchrony ^10,11^. Thus, it is possible that ASD risk gene mutations over-synchronize neural networks, leading to a reduction in network information encoding capability that disturbs cognitive performance^12–14^. Consistent with this theoretical framework, ASD animal models with increased E/I balance exhibit increased neuronal correlations, as well as deficits in social interaction^15,16^ and sensory discrimination^17^.

While lacking single neuron resolution, EEG variability analysis in humans has allowed the estimation of neural synchrony. As EEG provides an aggregate measure of neural activity-dependent extracellular electrical currents, lower EEG variability is indicative of greater neural synchrony. From a group of ASD subjects, those without detectable EEG epileptiform activity exhibit lower EEG variability and higher functional E/I ratios than typically developing children^18^. Further, lower EEG variability is associated with decreased accuracy on a facial recognition task in typically developing children^19^. Finally, a low-dose ketamine infusion in healthy adults, thought to increase E/I ratio, creates specific deficits in a spatial working memory task^20^. Together, these computational and experimental evidence, in both animal models and human subjects, indicate that E/I imbalance and neural synchrony could contribute to ASD network pathophysiology that ultimately results in behavioral disruptions.

Recently, two human genomic studies identified mutations in a X-linked gene, *NEXMIF* (also known as *KIDLIA, KIAA2022*, or *Xpn*), in several males who presented with ASD, intellectual disability, and other co-morbidities^21,22^. Since then, several additional studies have reported individuals with ASD with mutations or deletions in the *NEXMIF* gene^23–35^. *NEXMIF* is now recognized as an ASD-implicated gene in the Simons Foundation Autism Research Initiative (SFARI) database. *NEXMIF* is critical for proper neurite extension and migration in the developing mouse cortex^36^. In agreement with the E/I imbalance hypothesis, *NEXMIF* knockdown causes a 2-fold greater loss of GABAergic synapses compared to glutamatergic synapses in cultured neurons^37^. As *NEXMIF* is an X-linked gene, homozygous *NEXMIF* knockout in females is embryonically lethal. In male mice, however, genetic deletion of *NEXMIF* (NEXMIF KO) results in a variety of behavioral deficits, most notably reduced social interaction, impaired communication vocalizations, and increased self-grooming (indicative of repetitive behavior). While initially characterized in the cortex of developing mice, NEXMIF expression is highest in the hippocampus of adult mice^38,39^, and NEXMIF KO mice exhibit impaired spatial memory and contextual fear memory^37^. As hippocampal structure^40–44^ and function^45–48^ are often disrupted in individuals with ASD, NEXMIF KO animals allow mechanistic analysis of how *NEXMIF* deletion alters hippocampal networks.

To understand how ASD-implicated *NEXMIF* gene mutations alter hippocampal function at both the cellular and network levels, we performed wide-field calcium imaging of hundreds of individual CA1 neurons simultaneously in NEXMIF KO male mice and WT male littermates during locomotion. *NEXMIF* knockout causes few cellular-level effects, quantified by calcium event shape and frequency in individual neurons in KO and WT mice. However, we also characterized network effects of *NEXMIF* knockout using Pearson correlation and a network closeness centrality metric, and we found that loss of *NEXMIF* creates over-synchronization of the CA1 network during locomotion.

## Results

### NEXMIF wild-type (WT) and knockout (KO) mice exhibit similar locomotor behavior

NEXMIF expression is most prominent in the hippocampus, and NEXMIF KO mice exhibit severe learning and memory deficits^37,38^. To understand how NEXMIF contributes to hippocampal circuit functions, we characterized CA1 neural responses using calcium imaging while mice were head-fixed and navigating freely on a spherical treadmill (Figure 1A). Since it is very difficult for NEXMIF KO mice to perform hippocampal-dependent learning and memory tasks, we examined how *NEXMIF* knockout changes hippocampal circuity during locomotion, a fundamental component of spatial navigation.

**Figure 1.**
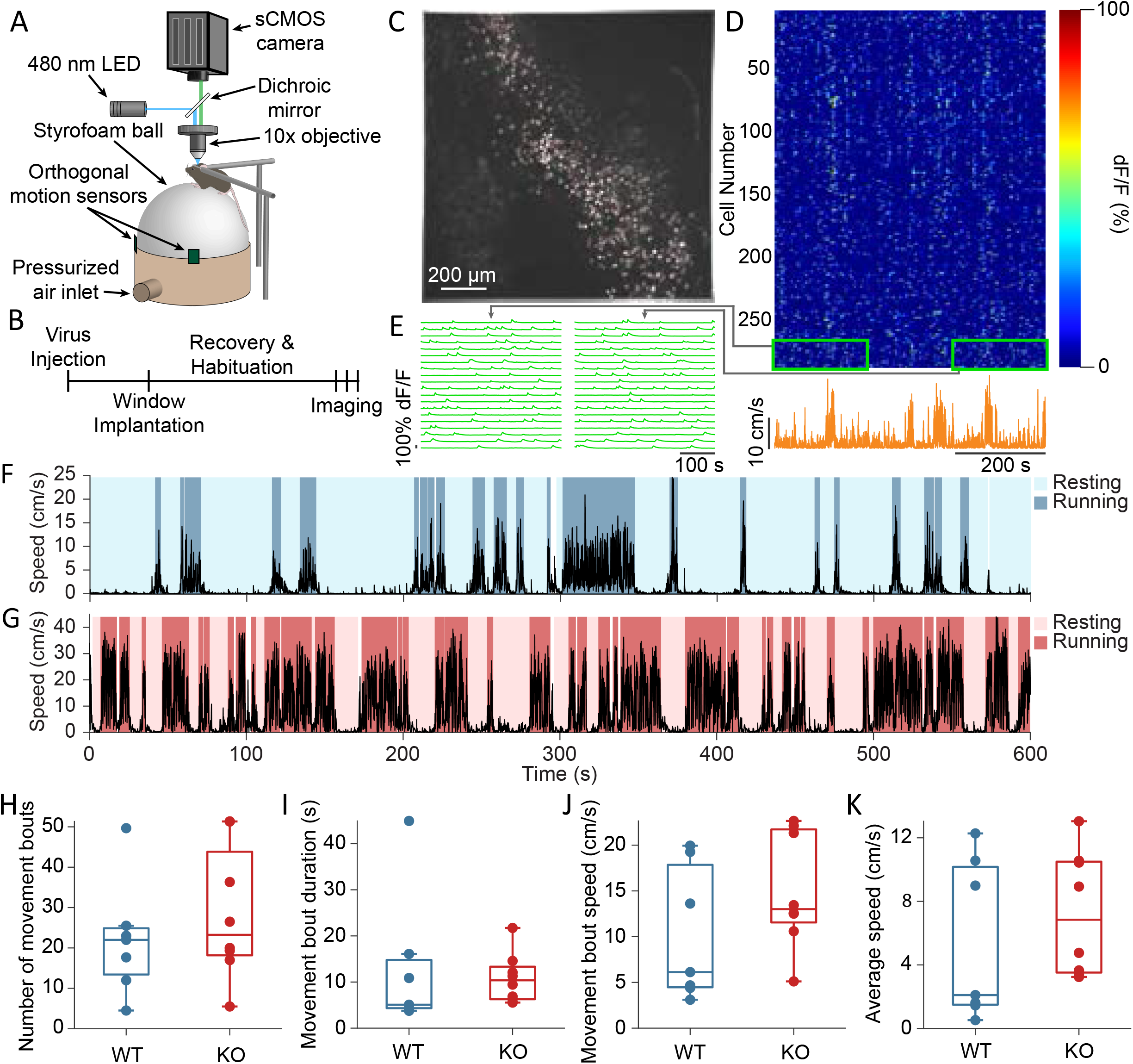
Experimental set-up and movement behavior. (A) Camera, microscope, and spherical treadmill set-up. (B) Experimental timeline. (C) Maximum-minus-minimum projection image across the entire video for one representative session. All selected ROIs are outlined in pink, with highlighted cells in D and E shown in green. (D) Heat map of GCaMP6f dF/F traces for the included ROIs shown in C during an example session (top) and animal’s movement speed from the same session (bottom). (E) Zoom-ins of the heat map regions outlined in green in D, showing the fluorescence traces for 20 representative cells from the beginning and the end of the imaging session. (F,G) Resting and running bouts identified in a representative (F) WT animal and (G) KO animal, overlaid on the movement speed from each animal. (H) Average number of movement bouts per session. p= 0.441, unpaired t-test. (I) Average movement bouts duration. p= 0.736, unpaired t-test. (J) Average speed during movement bouts. p= 0.188, unpaired t-test. (K) Average speed over the entire imaging session. p= 0.428, unpaired t-test. In H-K, each dot corresponds to an individual session. Box plots show median (middle line in box) and upper and lower quartiles (top and bottom edges of box, respectively). Whiskers show maximum and minimum values that are not outliers.

We first compared voluntary movement kinematics between male homozygous NEXMIF KO mice (n=8 mice) and male WT littermates (n=7 mice). “Resting” and “running” bouts were identified in each recording session based on movement speed (details in Methods, Figure 1F, G). WT and KO mice exhibited a similar number of running bouts (periods of continuous running) within each 10-minute session (Figure 1H, WT: 22.0±14.2 bouts, KO: 28.4±16.6 bouts, unpaired t-test, p=0.44, n=16 sessions from 7 WT mice & n=21 sessions from 8 KO mice), with similar running bout duration (Figure 1I, WT: 12.8±14.9 sec, KO: 10.9±5.43 sec, unpaired t-test, p= 0.73). As KO mice tend to run faster than WT mice during running bouts (Figure 1J, WT: 10.2±7.3 cm/s, KO: 15.04±6.33 cm/s, unpaired t-test, p= 0.18), average speed during the entire imaging session tends to be higher in KO mice, but these differences were not significant (Figure 1K, WT: 5.36±5.0 cm/s, KO: 7.25±3.92 cm/s, unpaired t-test, p= 0.43).

### Calcium event shape and frequency are undisturbed in NEXMIF KO mice

To examine the impact of *NEXMIF* knockout on both individual hippocampal neurons and the dorsal CA1 circuit, we performed calcium imaging from neurons expressing GCaMP6f, in both WT and KO animals (Figure 1B, C, D). Each mouse was recorded for 10 minutes per day during voluntary locomotion (Figure 1D, E). The recorded calcium fluorescence videos were motion corrected and individual cells were segmented^49^ (Figure 1C). A GCaMP6 fluorescence trace was then extracted for each cell and normalized to its peak fluorescence to account for variation in GCaMP6f expression between neurons. We then identified individual calcium events as described previously^50^ (Figure 2A).

**Figure 2.**
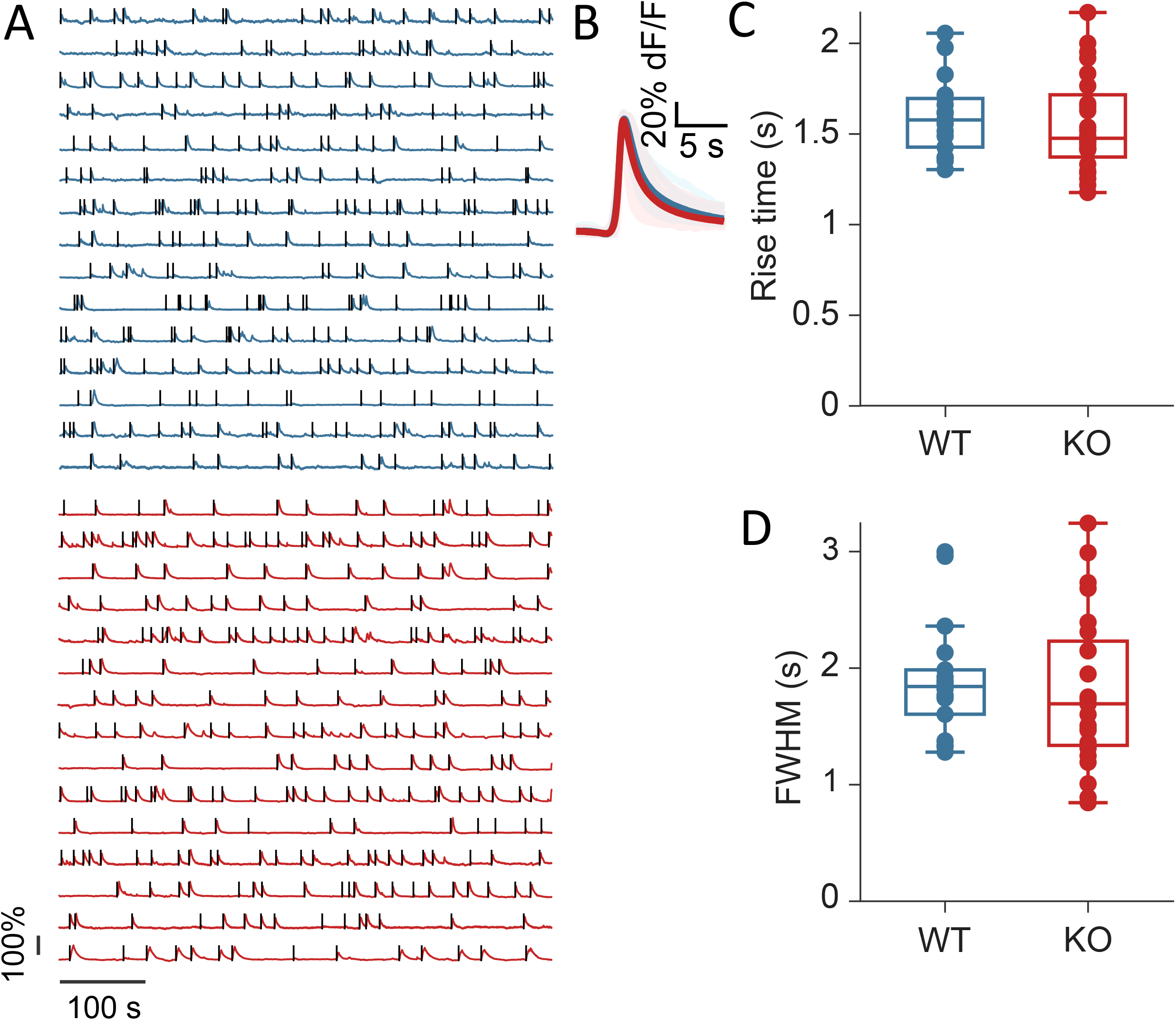
*NEXMIF* knockout does not change calcium event shape. (A) Example fluorescence traces from a WT animal (top) and a KO animal (bottom). 15 cells are shown from each mouse. Each detected calcium event is marked with a black line. (B) Average calcium event shape from all WT sessions (dark blue) and all KO sessions (dark red). Events were first averaged within each trace, then averaged across all cells in a session. Each session average is shown as a light line, and the population average is shown as the solid line. (C) Average calcium event rise time. Unpaired t-test, p= 0.60. (D) Average calcium event full width at half-maximum amplitude. Unpaired t-test, p= 0.66. In C and D, each dot corresponds to an individual session. Box plots show median (middle line in box) and upper and lower quartiles (top and bottom edges of box, respectively). Whiskers show maximum and minimum values that are not outliers.

We estimated neural activity using both the rise time and frequency of individual calcium events, as the rising phase of calcium events captures the sharp increases in intracellular calcium that are common during spike bursts^51^. We found that both calcium event rise time and calcium event frequency over the entire imaging session are similar between WT and KO mice (Figure 2B and C, n=18 sessions from 7 WT mice and 24 sessions from 8 KO mice, rise time WT: 1.59±0.22 ms, KO: 1.55±0.27 ms, unpaired t-test, p= 0.64; frequency WT: 2.1 ± 0.4 events/min, KO: 2.3 ± 0.5 events/min, unpaired t-test, p=0.22). Additionally, we used calcium event full width at half-maximum amplitude (FWHM) as a measure of calcium buffering capacity, because the duration of a calcium event captures overall intracellular calcium change^51,52^. We found that FWHM is also similar between WT and KO mice (Figure 2B-D, FWHM WT: 1.90±0.52 ms, KO: 1.81±0.66 ms, unpaired t-test, p= 0.65). Thus, *NEXMIF* knockout does not affect the overall activity or calcium buffering capacity of CA1 neurons.

### NEXMIF KO increases the fraction of movement-modulated neurons

Since CA1 neurons are known to increase their neural activity during locomotion^53,54^, we next compared calcium event rates during resting versus running. We found that calcium event rate across the entire neuron population increased during running in both WT and KO animals (Figure 3A, n= 12 sessions in 6 WT mice and n=20 sessions in 8 KO mice, WT rest: 1.71±0.51 events/min, WT run: 2.79±0.75 events/min, paired t-test with Bonferroni correction, p =7.76e-5; KO rest: 1.90±0.54 events/min, KO run: 2.93±0.75 events/min, paired t-test with Bonferroni correction, p= 5.08e-8). While KO mice have a slightly elevated event rate at the population level during both resting and running, this increase was not significant during either each locomotor state (Figure 3A, unpaired t-test with Bonferroni correction, WT vs KO rest: p= 0.32, WT vs KO run: p= 0.62).

**Figure 3.**
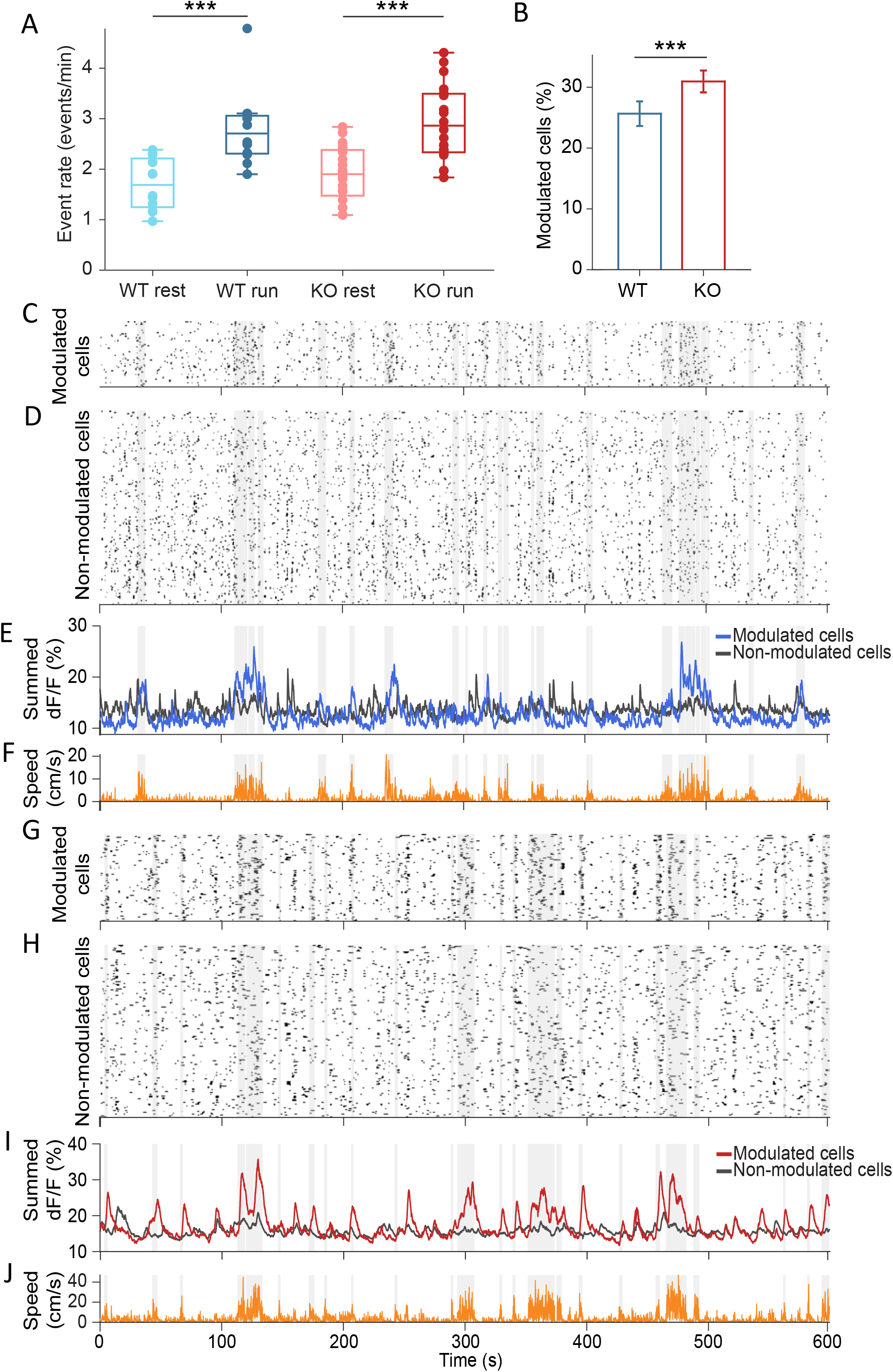
*NEXMIF* knockout increases percentage of movement-modulated cells. (A) Calcium event rate in WT mice during rest (light blue) versus run (dark blue), and in KO during rest (light red) versus run (dark red). Paired t-test with Bonferroni correction, WT rest vs WT run p= 7.76e-5; KO rest vs KO run: p= 5.08e-8. Unpaired t-test with Bonferroni correction, WT rest vs KO rest: p= 0.32; WT run vs KO run: p= 0.62. Each dot corresponds to an individual session. Box plots show median (middle line in box) and upper and lower quartiles (top and bottom edges of box, respectively). Whiskers show maximum and minimum values that are not outliers. ***p<0.00025. (B) Fraction of neurons that are movement modulated in WT (blue) versus KO (red) mice. Fisher’s exact test, p= 1.66e-4. Error bars represent 95% confidence intervals calculated using the standard error of proportions. ***p<0.001. (C-J) Example sessions from a (C-F) WT animal and (G-J) KO animal. Binarized calcium traces for all movement-modulated cells (C,G) and all non-movement-modulated cells (D,H) in the example sessions. (E,I) Summed normalized dF/F for movement-modulated cells (blue for WT and red for KO) and non-movement-modulated cells (black) in the session. (F,J) Corresponding movement speed (orange) for the session. All plots are overlaid with movement bouts in gray.

Since only some CA1 neurons are modulated by locomotion, we next examined whether the small increases in population calcium event rates observed in KO mice reflect a difference in the fraction of movement-modulated neurons. To determine whether a neuron is modulated by movement, we binarized the calcium event trace to represent total neural activity, with ones assigned to the entire rising phase of each calcium event and zeros everywhere else (Figure 3C, D, G, H). We then computed the difference in neural activity during running versus resting and compared it to a shuffled null distribution. In each shuffle, we circularly shifted each binarized calcium trace by a random temporal offset relative to movement and calculated the difference in activity between the running periods and resting periods. This procedure was repeated 1000 times to form the null distribution. A cell was deemed to be movement-modulated if the observed neural activity difference was greater than the 97.5^th^ percentile of the shuffled null distribution for that cell (Figure 3C, D, G, H). Using this analysis, we found that 31.0 ± 1.8% of neurons were movement-modulated in KO animals, significantly higher than the 25.7 ± 2.0% observed in WT (Figure 3B-J, n= 1805 cells from 6 WT mice and n= 2530 cells from 8 KO mice, Fisher’s exact test, p= 1.6e-4). The percentage of cells that were movement-modulated in each session did not depend on the time the animal spent running or the animal’s average speed during the session for either mouse group or behavioral condition (Figure S1). This increase in proportion of movement-modulated cells in KO mice suggests that *NEXMIF* knockout specifically increases behavioral responses of individual CA1 cells. As *NEXMIF* knockout increases E/I synaptic ratio of individual cells^37^, our results provide direct experimental evidence that increased E/I ratio by ASD risk gene mutation accordingly increases behaviorally-evoked responses, consistent with the observation that sensory stimuli lead to an over-activation of the hippocampus in individuals with ASD^48^.

### Neuronal correlation is increased in NEXMIF KO mice during both resting and running

Computational studies have shown that increased E/I ratio increases pairwise correlations amongst neuronal populations, thus decreasing network information coding capability^10–12^. Additionally, several animal models with deletions of ASD risk genes have increased neuronal correlations^16,17^. *NEXMIF* knockout increases E/I ratio, like many other ASD risk gene mutations, we next examined whether *NEXMIF* knockout influences CA1 neuronal correlations by calculating Pearson’s correlation between simultaneously recorded neuron pairs (Figure 4A, B). To account for variations in event rate, we determined whether the measured correlation between each neuron pair was significantly greater than chance observation given the event rates of the neurons in the pair. To estimate chance observations, we shifted the binarized traces of two neurons relative to one another with a random time lag and obtained a shuffled Pearson correlation coefficient. We repeated this shuffling procedure 2000 times to create a shuffled null distribution. If the observed correlation coefficient was greater than the 95^th^ percentile of the shuffled null distribution, the neuron pair was deemed significantly correlated (correlated pair). If the observed correlation coefficient was below the 95^th^ percentile of the shuffled distribution, the correlation was deemed non-significant (random pair). We found KO animals contained more correlated cell pairs compared to WT mice, during both resting and running conditions (Figure 4C, Fisher’s exact test, WT rest vs KO rest: p= 7.7e-32; WT run vs KO run: p= 3.4e-127). The number of correlated pairs dropped during running in both WT and KO mice (Figure 4C, WT rest: 8.12±0.10%, WT run: 6.55±0.09%, Fisher’s exact test, p= 2.1e-121; KO rest: 8.93±0.09%, KO run: 8.09±0.09%, Fisher’s exact test, p= 1.76e-37). These results indicate that locomotion desynchronizes the CA1 network, and *NEXMIF* knockout leads to abnormally elevated synchronization regardless of behavioral conditions.

**Figure 4.**
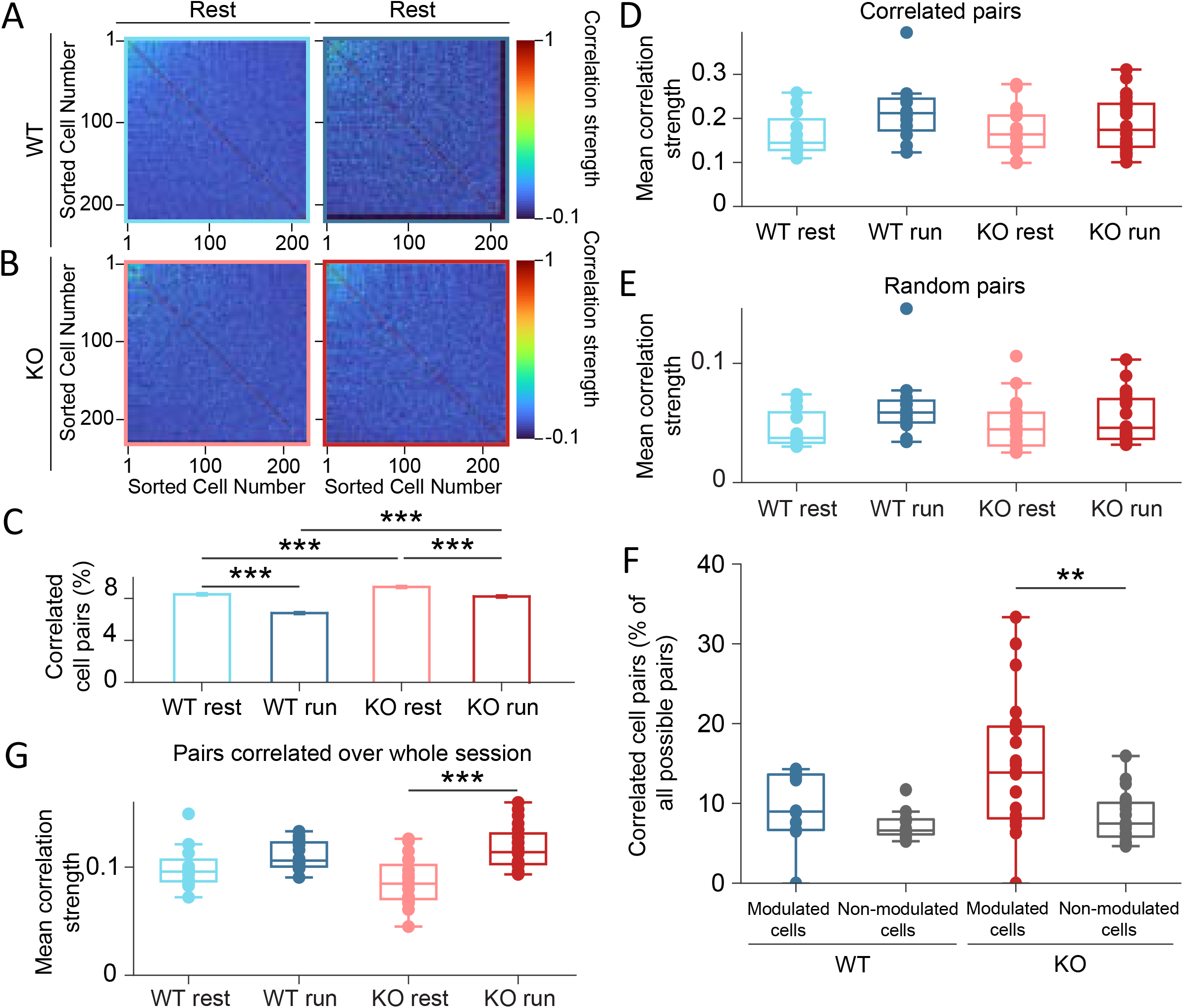
Pairwise correlation increases during running in NEXMIF KO mice. (A) Correlation matrices of average pairwise Pearson correlation coefficient during resting (left) and running (right) periods for all neurons from an example WT animal. Within each matrix, neurons are sorted such that correlated cell pairs are shown in the top left corner. (B) Same as in A, for an example KO animal. (C) Fraction of neuron pairs that are correlated in WT mice during rest (light blue) and run (dark blue), and in KO during rest (light red) and run (dark red). Fisher’s exact test, WT rest vs KO rest: p= 7.7e-32, WT run vs KO run: p= 3.4e-127, WT rest vs WT run: p= 2.1e-121, KO rest vs KO run: p= 1.8e-37. Error bars represent 95% confidence intervals calculated using the standard error of proportions. ***p<0.001 (D) Average Pearson correlation coefficients during resting or running for cell pairs correlated in that behavioral condition. Paired t-test with Bonferroni correction, WT rest vs WT run: p = 0.12; KO rest vs KO run p= 0.50, unpaired t-test with Bonferroni correction, WT rest vs KO rest: p= 0.70, WT run vs KO run: p= 0.22. (E) Same as D, for random cell pairs. Paired t-test with Bonferroni correction, WT rest vs WT run: p= 0.12, KO rest vs KO run: p= 0.53; unpaired t-test with Bonferroni correction, WT rest vs KO rest: p=062, WT run vs KO run: p= 0.26. (F) Fraction of movement-modulated or non-modulated cell pairs that are significantly correlated for all sessions. Paired t-test with Bonferroni correction, WT modulated vs WT non-modulated: p= 0.09, KO modulated vs KO non-modulated: p= 6.5e-4, unpaired t-test with Bonferroni correction, WT modulated vs KO modulated: p= 0.06, WT non-modulated vs KO non-modulated: p= 0.30. (G) Average Pearson correlation coefficients during resting or running of cell pairs correlated over entire session. Paired t-test with Bonferroni correction, WT rest vs WT run: p=0.18; KO rest vs KO run p= 1.2e-5, unpaired t-test with Bonferroni correction, WT rest vs KO rest: p= 0.09, WT run vs KO run: p= 0.29. **p<0.00125, ***p<0.000125. In D-G, each dot corresponds to an individual session. Box plots show median (middle line in box) and upper and lower quartiles (top and bottom edges of box, respectively). Whiskers show maximum and minimum values that are not outliers.

We then evaluated the strength of correlation by comparing correlation coefficients. We found that correlation coefficients between correlated pairs were similar between WT and KO mice during both resting and running (Figure 4D, n= 12 sessions in 6 WT mice and n=20 sessions in 8 KO mice, correlated pairs: WT rest: 0.16±0.05, WT run: 0.22±0.07, KO rest: 0.17±0.05, KO run: 0.19±0.06, paired t-test with Bonferroni correction, WT rest vs WT run: p= 0.12, KO rest vs KO run: p= 0.50; unpaired t-test with Bonferroni correction, WT rest vs KO rest: p= 0.70, WT run vs KO run: p= 0.22). The correlation coefficients of random pairs were also similar between WT and KO during both behavioral conditions (Figure 4E, n= 12 sessions in 6 WT mice and n=20 sessions in 8 KO mice, random pairs: WT rest: 0.05±0.02, WT run: 0.06±0.03, KO rest: 0.05±0.02, KO run: 0.05±0.02, paired t-test with Bonferroni correction, WT rest vs WT run: p= 0.12, KO rest vs KO run: p= 0.53; unpaired t-test with Bonferroni correction, WT rest vs KO rest: p= 0.62, WT run vs KO run: p= 0.26). Thus, *NEXMIF* knockout increases functional network connectivity regardless of behavioral condition by synchronizing more neurons, without altering the strength of the connectivity.

### In KO animals, a larger fraction of movement-modulated cells are correlated during running

To directly compare how CA1 network correlation strength differs between the two behavioral states, we identified correlated pairs that are session-relevant by calculating pairwise Pearson correlations over the entire recording session. In WT mice, the correlation strengths of these session-relevant cells were the same between resting and running, indicating that when an animal switches between the two behavioral states, CA1 network connectivity remains stable (Figure 4G, WT rest: 0.10, WT run: 0.11 paired t-test with Bonferroni correction, p=0.18). In KO mice, however, correlation strength among session-relevant cells is significantly higher during running than resting, indicating that with loss of NEXMIF, functional connectivity in the CA1 network is predominately driven by correlated activity during running (Figure 4G, KO rest: 0.09, KO run: 0.12, paired t-test with Bonferroni correction, p=1.2e-5; unpaired t-test with Bonferroni correction, WT rest vs KO rest: p=0.09, WT run vs KO run: p=0.29). This result indicates that *NEXMIF* knockout leads to over-synchronization during running.

As locomotion evoked responses from a larger fraction of cells in KO animals (Figure 3), we next examined whether functional connectivity differs between populations of movement-modulated cells versus non-movement-modulated cells. We first identified correlated pairs among movement-modulated cells versus non-movement-modulated cells. In WT animals, similar fractions of correlated cells were observed amongst modulated cells and non-modulated cells (Figure 4F, modulated cell pairs: 9.5±4.4%, non-modulated cell pairs: 7.3±1.8%, paired t-test with Bonferroni correction, p= 0.09). However, in KO animals, a strikingly larger fraction of movement-modulated cells were correlated (14.8±8.6%) compared to non-movement-modulated cells (8.3±3.1%, Figure 4F, paired t-test with Bonferroni correction, p= 6.5e-4). Correlation strength was similar for correlated pairs amongst movement-modulated cells versus non-movement-modulated cells in both WT and KO animals (WT modulated 0.10±0.04 vs non-modulated 0.10±0.04: p=0.26; KO modulated 0.09±0.04 vs non-modulated 0.08±0.03: p= 0.16, paired t-test with Bonferroni correction). Thus, *NEXMIF* knockout not only increases the fraction of neurons that respond to movement, but also increases the functional connectivity amongst movement-modulated cells.

### CA1 network connectivity is exaggerated during locomotion in NEXMIF KO

To further characterize functional connectivity within the CA1 network, we next created network maps using the correlated cell pairs during either resting or running. Each cell is a node in the network, and a correlated cell pair is connected by an edge between those two nodes. As fluorescence imaging allowed us to visualize the anatomical relationship between recorded neurons, we first arranged the network using the anatomical position of each cell (Figure 5A). We did not observe any obvious patterns in the closeness centrality within CA1 networks in the anatomical maps. Consequently, to better visualize network connectivity, we arranged each map as a force-directed graph where cells are positioned based on the strength of their functional connectivity with other cells rather than absolute anatomical location (Figure 5B). In WT force-directed maps, cells were more tightly clustered, indicating higher connectivity, during resting versus running. However, KO force-directed maps showed similar patterns of clustering across resting and running (Figure 5B).

**Figure 5.**
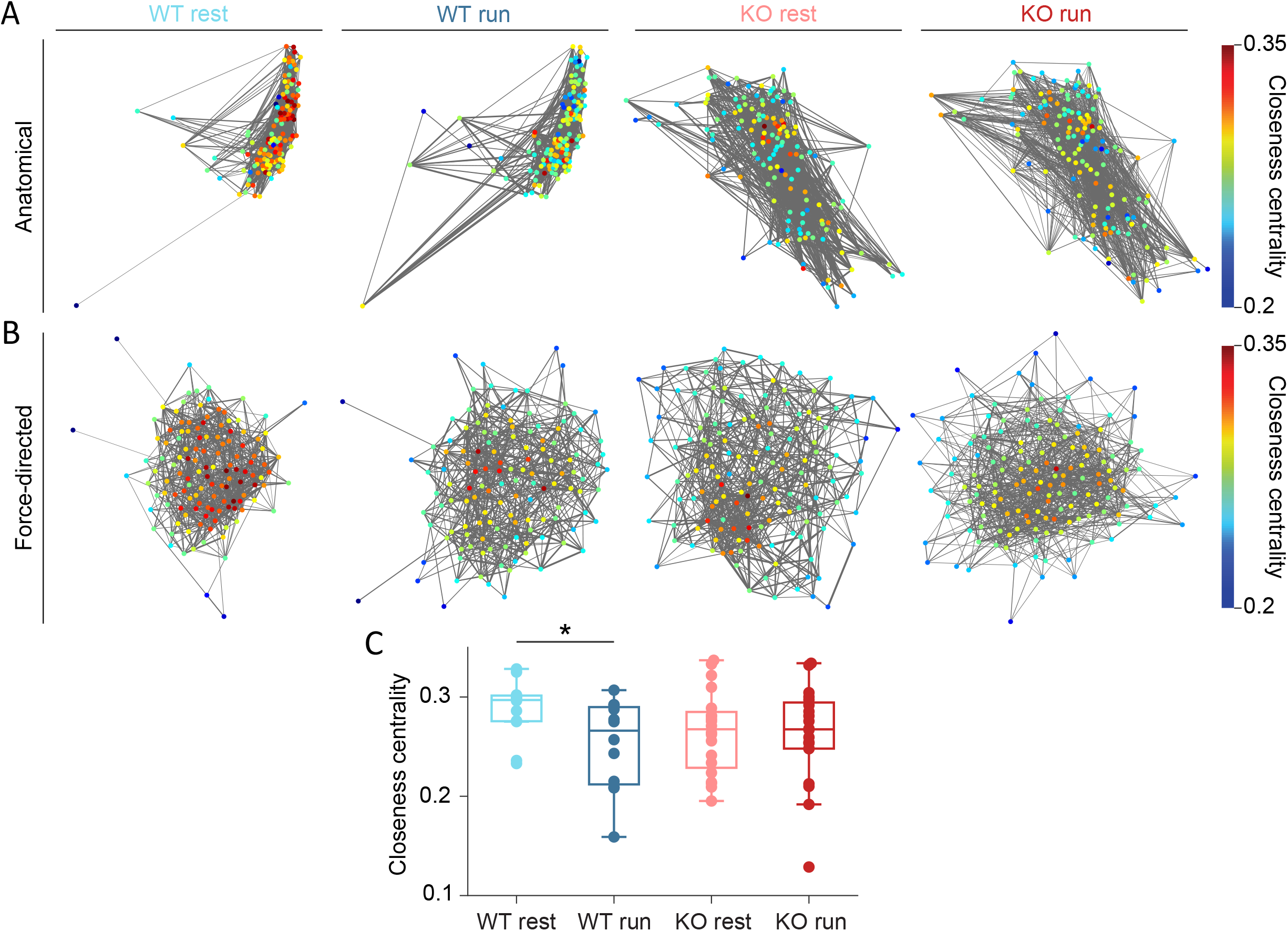
*NEXMIF* knockout increases functional connectivity of the CA1 network. (A) Anatomical network maps during resting and running from example WT and KO animals. Each cell is color coded based on its closeness centrality measure. Correlated cell pairs (identified in each behavioral condition separately) are connected by an edge. Edge width represents normalized correlation coefficient. (B) Same as A, shown as a force-directed graph. (C) Average closeness centrality in WT mice during rest (light blue) versus run (dark blue), and in KO during rest (light red) versus run (dark red). Paired t-test with Bonferroni correction, WT rest vs WT run: p= 0.01; KO rest vs KO run p= 0.76, unpaired t-test with Bonferroni correction, WT rest vs KO rest: p= 0.1, WT run vs KO run: p= 0.56. Each dot corresponds to an individual session. Box plots show median (middle line in box) and upper and lower quartiles (top and bottom edges of box, respectively). Whiskers show maximum and minimum values that are not outliers. *p<0.0125

Thus, to quantify the connectivity of each network, we calculated its average closeness centrality, which takes both number of connections and edge strength (correlation coefficient) into account (Methods). Briefly, a greater closeness centrality value for a neuron indicates that the neuron is connected, both directly and indirectly, to a greater number of nodes in the network. In accordance with our observations of the force-directed maps, we found that closeness centrality decreases in WT mice during running compared to resting, indicating that the CA1 network is less interconnected, and thus desynchronized, during locomotion (Figure 5C, WT rest: 28.8, WT run: 25.2, paired t-test with Bonferroni correction, p=0.01). This overall network-level desynchronization is consistent with our observations that fewer cell pairs were correlated during locomotion and that correlated cell pairs showed similar correlation strength in WT mice (Figure 4C, D). In contrast, in KO mice, closeness centrality did not change between running and resting, indicating that the KO network fails to desynchronize during locomotion (Figure 5C, KO rest: 26.4, KO run:26.2, paired t-test with Bonferroni correction, p=0.76; unpaired t-test with Bonferroni correction, WT rest vs KO rest: p=0.1, WT run vs KO run: p=0.56). In KO mice, the fraction of correlated cells is higher than WT in both behavioral conditions, and amongst cell pairs correlated amongst the whole session, correlation strength increased during running (Figure 4C, G). The interaction of these two effects, which occur at the level of cell pairs, could contribute to the overall lack of change in functional connectivity at the network level. Together, these results demonstrate that while the WT CA1 network desynchronizes during locomotion, the NEXMIF KO CA1 network fails to do so.

## Discussion

In this study, we examined how loss of *NEXMIF*, an ASD risk gene, influences individual CA1 neuron’s response and CA1 network connectivity during locomotion. As we used single-cell resolution calcium imaging during voluntary navigation to interrogate the hippocampal CA1 network, we are able to compare the patterns of neural activation during quiescent immobility versus active locomotion. We found that spontaneous calcium event rate is similar between WT and KO mice, but a larger percentage of CA1 neurons are activated during movement in KO mice. Furthermore, a greater fraction of neurons within the CA1 network are correlated in KO animals, and movement-responsive cells are more correlated in KO animal than in WT animals. Finally, we also found that the KO network is overly connected during locomotion specifically. Overall, our results demonstrate that *NEXMIF* knockout leads to increased behaviorally-evoked responses and elevated network synchronization, both of which could contribute to the disruption of CA1 network coding ability during behavior.

As increased cellular and synaptic level E/I ratio in ASD can lead to increased neuronal excitability, ASD is often co-morbid with epilepsy, particularly in individuals with *NEXMIF* mutations^3,35^. We did not observe differences in basal calcium event rate in NEXMIF KO mice, which indicates the presence of homeostatic mechanisms that counteract the increased synaptic E/I ratio shown previously by Gilbert et al.^37,55^. However, we detected significantly more neurons that selectively increased their activity during movement in KO animals, providing direct evidence that in *NEXMIF* knockout, elevated synaptic E/I ratio does not translate to a broad increase in spontaneous neuronal excitability, but rather a selective increase in responding during relevant behavior. This behavioral state-specific increase in neuronal excitability in the CA1 could be due to a global increase in synaptic inputs to CA1 during movement, in which increased E/I synaptic ratio leads to greater excitatory drive to CA1 neurons. However, it is also possible that the observed excitability increase reflects movement-dependent changes in intrinsic biophysical neuronal properties, in addition to altered synaptic inputs.

Another leading hypothesis of ASD pathophysiology argues for overconnectivity within local brain regions and underconnectivity between interconnected brain regions, supported by several exciting human studies^56–62^. We observed increased fractions of correlated CA1 neuron pairs in KO animals during both immobility and active locomotion. Additionally, in session-relevant cell pairs from KO mice, correlation strength increases during running. Most strikingly, we also detected a selective increase in the number of correlated cell pairs within movement-relevant cells, and no difference in closeness centrality during running. Each of these observations is consistent with increased functional connectivity within the CA1 circuit of NEXMIF KO mice during locomotion, supporting the local overconnectivity hypothesis. Elevated E/I synaptic ratio could contribute to this increased functional connectivity^10,11^, but again, we cannot rule out the possibility that *NEXMIF* knockout also changes intrinsic biophysical properties that lead to the observed over-synchronization. Further work is needed to better understand the exact mechanisms by which *NEXMIF* alters both cellular biophysical properties such as ion channel expression and functional connectivity between the hippocampus and its cortical targets.

While the fractions of correlated pairs increased in KO animals, the correlation strength of these pairs remained constant across behavioral conditions in WT and KO animals, indicating that functional connectivity strength between neurons is not sensitive to behavioral state or *NEXMIF* knockout. Interestingly, the number of correlated cells decreases in WT animals during locomotion, and WT animals exhibit fewer correlated cell pairs amongst locomotion-relevant cells. Additionally, closeness centrality of the WT network decreases during running, confirming decreased functional connectivity in the WT CA1 network during movement. Desynchronization of the neuronal population would lead to increased information encoding capability, consistent with the idea that CA1 network encodes relevant information during movement^63^. As locomotion is a fundamental component of spatial navigation and memory, this dynamic change in information coding capability would allow for flexible and efficient encoding of a WT animal’s current environment for spatial memory.

In NEXMIF KO animals, however, a larger number of cell pairs are significantly correlated in both behavioral conditions, and cell pairs that are correlated over the entire session are dominated by strong correlations during running. Additionally, a larger fraction of locomotion-relevant cells are highly correlated, and closeness centrality of the CA1 network does not decrease during movement. Taken together, these observations illustrate that *NEXMIF* knockout creates exaggerated network synchrony and thus reduces information encoding capacity, particularly during locomotion. Although a larger population of cells responds during movement in KO animals, the abnormal synchrony amongst these cells and others prohibits the heterogeneity of CA1 network response, illustrated by closeness centrality, that is seen in WT mice during locomotion. The increased synaptic E/I ratio of NEXMIF KO neurons and our finding of overexcitation upon movement in the CA1 circuit of KO mice could account for the similarity we observe in population activity. The higher percentage of movement-modulated cells observed in KO mice could also reflect a compensatory mechanism in the CA1, to homeostatically increase information encoding capability throughout development. Alternatively, this higher percentage could be due to the increased number of correlated cells, as these correlations could arise from common inputs to these cell pairs that are activated upon movement. Overall, our observations of increased connectivity indicate a reduced ability to process spatial information and spatial encoding that could lead to the impaired spatial memory and contextual fear memory observed in NEXMIF KO mice^37^.

## Materials and Methods

### Animal Surgery and Recovery

All animal procedures were approved by the Boston University Institutional Animal Care and Use Committee. 8 homozygous *KIAA2022* knockout (maintained on a C57Bl/6 genetic background) male mice and 7 wild-type (WT) male littermates were used in this study^37^. Mice were 7-34 weeks old at the start of experiments. Animals first underwent stereotaxic viral injection surgery, targeting the hippocampus (anterior/posterior: -2.0mm, medial/lateral: +1.4mm, dorsal/ventral: -1.6mm from bregma). Mice were injected with 500-750nL of AAV9-synapsin-GCaMP6f.WPRE.SV40 virus, obtained from the University of Pennsylvania Vector Core (titer ∼6e12 GC/mL). Injections were performed with a blunt 33-gauge stainless steel needle (NF33BL-2, World Precision Instruments) and a 10 μL microinjection syringe (Nanofil, World Precision Instruments), using a microinjector pump (UMP3 UltraMicroPump, World Precision Instruments). The needle was lowered over 1 min and remained in place for 1 min before infusion. The rate of infusion was 50 nL/min. After infusion, the needle remained in place for 7-10 min before being withdrawn over 1 min. The skin was then sutured closed with a tissue adhesive (Vetbond, 3M). After complete recovery (7+ days after virus injection), animals underwent a second surgery to implant a sterilized custom imaging cannula (outer diameter: 3.17mm, inner diameter: 2.36mm, height: 2mm). The imaging cannula was fitted with a circular coverslip (size 0, outer diameter: 3 mm, Deckgläser Cover Glasses, Warner Instruments), adhered to the bottom using a UV-curable optical adhesive (Norland Optical Adhesive 60, P/N 6001, Norland Products). During surgery, an approximately 3.2mm craniotomy was created (centered at anterior/posterior: -2.0mm, medial/lateral: +1.7mm) and the cortical tissue overlaying the hippocampus was aspirated away to expose the corpus callosum. The corpus callosum was then thinned until the underlying CA1 became visible. The imaging cannula was then tightly fit over the hippocampus and sealed in place using a surgical silicone adhesive (Kwik-Sil, World Precision Instruments). The imaging window was secured in place using bone adhesive (C&B Metabond, Parkell) and dental cement (Stoelting). A custom aluminum head-plate was also affixed to the skull anterior to the imaging window. Analgesic was provided for at least 48 hours after each surgery, and mice were single-housed after window implantation surgery to prevent damage to the head-plate and imaging window.

### Animal Habituation, Calcium Imaging, and Movement Data Acquisition

After complete recovery from window implantation surgery (7+ days), animals were habituated to experimenter handling and head fixation on the spherical treadmill. Each animal was habituated to running on the spherical treadmill while head-fixed for at least 3 days prior to the first recording day. During each recording session, animals were positioned under a custom wide-field microscope and allowed to run freely on the spherical treadmill. The spherical treadmills consisted of a three-dimensional printed plastic housing and a Styrofoam ball supported by air^64^. The imaging microscope was equipped with a scientific complementary metal oxide semiconductor (sCMOS) camera (ORCA-Flash4.0 LT Digital CMOS camera C11440-42U, Hamamatsu) and a 10× 0.28 M Plan Apo objective (Mitutoyo). GCaMP6f excitation was accomplished with a 5 W light emitting diode (M470L4, ThorLabs). The microscope included an excitation filter (No. FF01-468/553-25, Semrock), a dichroic mirror (No. FF493/574-Di01-25×36, Semrock), and an emission filter (No. FF01-512/630-25, Semrock). The imaging field of view was 1.343 × 1.343mm (1024 × 1024 pixels). Image acquisition was performed using HC Image Live (Hamamatsu), and images were stored offline as multi-page tagged image files (TIFs) for further analysis.

Each animal underwent three 10-minute recording sessions, one per day, every other day over 5 days (Figure 1B). A total of 21 recording sessions were collected from 8 KO mice and 16 sessions were collected from 7 WT mice. In 24 recording sessions (from 4 WT mice and 8 KO mice), a custom MATLAB script was used to trigger image frame capture at 20 Hz and to synchronize image acquisition with movement tracking. Digital transistor-transistor logic (TTL) pulses were delivered to the camera via a common input/output interface (No. USB-6259, National Instruments), and TTL pulses were also recorded using a commercial system (RZ5D, Tucker Davis Technologies). Motion data was collected using a modified ViRMEn system^65^. Movement was tracked using two computer universal serial bus mouse sensors affixed to the plastic housing at the equator of the Styrofoam ball, 78° apart. The mouse sensors’ x- and y-surface displacement data were acquired at 100 Hz on a separate computer, and a multi-threaded Python script was used to send packaged <dx,dy> data to the image acquisition computer via a RS232 serial link. Packaged motion data was recorded on the image acquisition computer using a modified ViRMEn MATLAB script and synchronized to each acquired imaging frame.

In the remaining 13 sessions (from 4 WT mice and 2 KO mice), image acquisition was triggered using a Teensy microcontroller system^66^, and experiments were performed using an identical spherical treadmill. Digital pulses were sent from a Teensy 3.2 (TEENSY32, PJRC) to the sCMOS camera via SMA connectors and coaxial cables to trigger frame capture at 20 Hz. TTL pulses were recorded using the same TDT commercial system. Movement was tracked using two computer mouse sensors (ADNS-9800 laser motion sensors, Tindie) affixed to the plastic housing at the equator of the Styrofoam ball, about 75 degrees apart. The x- and y-surface displacement was collected by the Teensy at 20Hz and sent to the image acquisition computer via a standard USB-microUSB cable.

### Movement Analysis

As both movement data acquisition systems collect the same numerical data, linear velocity can be calculated the same way for all sessions. Linear velocity in perpendicular X and Y directions was calculated as:

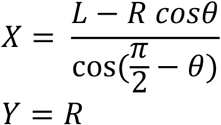

where L is the vertical reading from the left sensor, R is the vertical reading from the right sensor, and θ is the angle between the sensors. Linear velocity V was then calculated as:

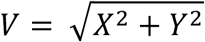

Linear velocity values were then interpolated at 20 Hz.

To identify sustained periods of movement with high linear velocity (running bouts), we used a Fuzzy logic-based thresholding algorithm. We first assigned each velocity data point a Fuzzy membership value using a sigmoidal membership function F:

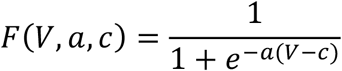

where the threshold c is the 20th percentile of the velocity of that session or 5 cm/sec, whichever is higher. a is set at 0.8. The resulting velocity trace was then smoothed using a moving average filter of 1.5 seconds. Next, the smoothed trace was thresholded at 10% of its maximum value. Time periods with velocity higher than this threshold that were at least 2 sec long were considered high velocity periods (“running”). Time periods with velocity lower than this threshold that were at least 2 sec long were considered low velocity periods (“resting”). Periods that do not satisfy either of these requirements were not considered for locomotion analysis (Figure 1F, G). Recording sessions in which the mouse spent less than 60 seconds (10% of the session) in one behavioral state and sessions with less than 5 running bouts were not included for locomotion-related analysis (4 sessions from 2 WT mice and 1 session from 1 KO mouse).

### Calcium Imaging Video Motion Correction

Videos were first motion corrected using a custom Python script as described previously^67^. For each imaging session, we first generated a reference image by calculating the mean value of each pixel across the first 2,047 frames. We then performed a series of contrast-enhancing procedures to highlight image features as follows. We used a Gaussian filter (python scipy package, ndimage.Gaussian_filter, sigma=50) to remove the low-frequency component, which represents the potential non-uniform background. We then captured the edges of the high-intensity area by calculating the differences between two Gaussian filtered images (sigma=2 and 1). To obtain the edge-enhanced image, the edges were multiplied by 100 and added back to the first filtered image (sigma=2). Finally, to prevent a potential overall intensity shift caused by photobleaching, we normalized the intensity of each image by subtracting the mean intensity of the image from each pixel and dividing by the standard deviation of the intensity. We then calculated the cross-correlations between the processed reference image and each processed image frame, and obtained the displacement between the peak of the coefficient and the center of the image. We then applied a horizontal shift, opposite to the displacement, to the original frame to finalize the motion correction.

### Cell Segmentation

From the motion corrected video, a projection image was generated across all frames by subtracting the minimum fluorescence from the maximum fluorescence of each pixel (max-min projection image), and regions of interest (ROIs) corresponding to fluorescent cell bodies were automatically identified in the max-min projection image using a deep-learning algorithm based on U-Net^68–70^. We first trained the deep-learning algorithm with manually curated data, containing the datasets reported in our previous studies^49,50^. For each training dataset, a max-min projection image was calculated as described above. We then divided the projection images and their corresponding ROI masks into small patches of 32×32 pixels as our training dataset. We also normalized each patch by shifting its mean intensity to zero and dividing the intensity of each pixel by the standard deviation of the patch intensity. During training, each pixel was further augmented by randomly flipping vertically and/or horizontally, and rotating 90, 180, or 270 degrees.

To segment ROIs for the datasets in this study, the max-min projection image for each dataset was divided into 32×32 patches with 50% of each patch overlapping with its neighboring patches. Each patch was normalized as described above. As a result, each pixel was inferred four times, and we averaged the results from four inferences as the prediction score. The connected pixels with a predication score >0.5 were segmented as a potential ROI, and the set of segmentations was further refined with watershed transformation to obtain the ROIs representing single neurons. ROIs were then overlaid on the max-min projection image and manually inspected. ROIs that were identified by the machine learning algorithm but were not identified as a cell by the observer were manually removed. ROIs were then manually added to select cells that the machine learning algorithm did not properly identify. ROIs were added as a circle with a radius of 6 pixels (7.8 μm) based on morphology present in the max-min projection image, using the previously-reported semi-automated custom MATLAB software called SemiSeg (github.com/HanLabBU/SemiSeg)^71^.

### GCaMP6f Fluorescence Trace Extraction and Normalization

We obtained the GCaMP6f fluorescence for each cell as the average fluorescence intensity across all pixels in that ROI. We then subtracted background fluorescence from each ROI, where the background fluorescence is the average pixel intensity across the pixels located within a ring centered at the corresponding cell ROI with an outer radius of 50 pixels and an inner radius of 15 pixels. The areas corresponding to other cell ROIs were excluded from this background ROI. Because the motion correction procedure introduces strips with high pixel intensities along the edges of the max-min projection image, 25 pixels along each edge of the image were also excluded from the calculation of background fluorescence. The resulting fluorescence trace for each cell was then interpolated at 20 Hz, linearly detrended (MATLAB function detrend), and normalized between 0 and 1. All traces were then manually inspected. Traces with large artifacts were removed.

### Calcium Event Detection

Onsets of calcium events were identified in each fluorescence trace similarly to previous descriptions^50,71^. Briefly, we first applied a moving average filter of 1s to smoothen each trace and calculated the spectrogram (MATLAB chronux, mtspecgramc with tapers = [2, 3] and window = [1, 0.05]), and averaged the power below 2 Hz. We then calculated the change in power at each time point (powerdiff) and identified outliers (3 median absolute deviations away from the median power) in powerdiff (MATLAB function isoutlier) to detect all significant changes in trace power. When multiple outliers occurred at consecutive time points, they were classified as a potential calcium event. We then calculated the rise time and amplitude (the difference in fluorescence value between the peak and the event onset) for each potential event and used an iterative process to include only true events and exclude incorrect potential events. Within each iteration, an amplitude threshold was calculated for each potential event (iteration 1: 7 standard deviations of the trace in the 10 seconds (200 data points) prior to calcium event onset). Potential events with a rise time greater than 150 ms (3 data points) and an amplitude above the calculated threshold were marked as correctly identified events for analysis. All the data points corresponding to these events (from beginning of event rise to end of event fall) were removed prior to the next iteration. We then repeated this process by re-calculating the amplitude threshold for the remaining potential events and again marking correctly identified events for analysis using the same criterion for rise time and the new amplitude threshold. For each successive iteration, the amplitude threshold was decreased by 40% and the duration to inspect prior to calcium event onset was increased by 75%. The iterative process stops once no events are marked as correctly identified events. This iterative method is more robust in capturing events that occur close together, while only minimally increasing identification of false positives. The preceding event in a sequence will incorrectly bias the standard deviation of the trace in the window prior to a following event in the sequence, and this bias is removed when the preceding event is not included in the window prior to the event onset. All traces were then manually inspected to confirm event detection accuracy.

### Calcium Event Parameter and Event Rate Analysis

For each detected calcium event, the rise time is defined as the duration from the calcium event onset, t_on,_ to its peak t_peak_. The full width at half-maximum amplitude (FWHM) was identified as the duration between two points t_a_ and t_b_ such that:

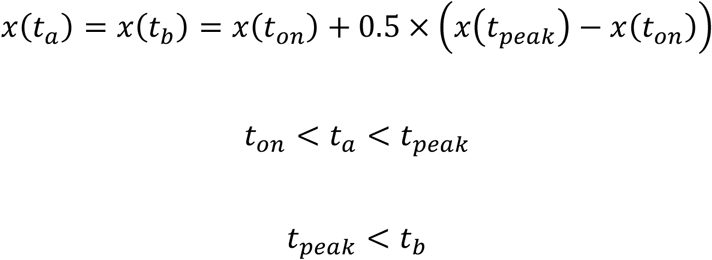

where *x*(*t*) is the fluorescence trace. Total event rate was calculated across the entirety of each trace, counting each identified calcium event as one event. Event rate during either running or resting was calculated by counting the number of calcium events in all bouts of the relevant behavioral condition and dividing by the total time that the mouse spent in that behavioral condition.

### Determination of Movement-Modulated Cells

To determine movement-modulated cells, we binarized each fluorescence trace by assigning ones to the entire rising phase (t_on_ to t_peak_) of each calcium event and zeros to the rest of the trace. We then concatenated all of the resting or running bouts separately, and summated the binarized trace among each concatenated period (“total activity”). We then subtracted the total activity during resting from the total activity during running to create an activity metric A. The calculation can be summarized as:

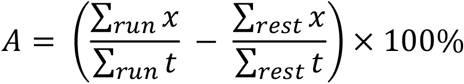

Next, we created a shuffled distribution of the activity metric for each cell by randomly circularly shifting the binarized trace relative to the movement trace 1000 times and calculating *A* for each shuffle. If the true (non-shifted) *A* for a neuron was greater than the 97.5th percentile of the shuffled distribution, the cell was considered movement-modulated. Cells that do not meet this criterion were considered non-movement-modulated.

### Pairwise Pearson Correlation Analysis

For pairwise correlation analysis, we calculated the Pearson’s correlation coefficient between the binarized traces for each pair of neurons. Only neuron pairs that were at least 20 pixels (26.2 μm) apart were included in all correlation analysis to eliminate potential fluorescence cross-contamination. We calculated pairwise correlation during resting alone, during running alone or during the entire duration of the session. To calculate pairwise correlation during resting alone or running alone, we concatenated the calcium activity during all resting or running periods. To calculate pairwise correlation during the entire duration of the session, we used the full length of the calcium traces for each cell pair.

To determine whether the correlation coefficient for each cell pair was above chance level for each behavioral condition, we created a shuffled distribution of correlation by randomly circularly shifting one trace relative to the other trace 2000 times and calculating the Pearson’s correlation coefficient for each shuffle. If the true (non-shifted) Pearson’s correlation for a pair of neurons was greater than the 95^th^ percentile of the shuffled distribution, the cell pair was considered correlated. Positive correlation coefficients between neuron pairs that were not greater than the 95^th^ percentile were considered random. Negative correlations were not included in any analyses due to the sparseness of GCaMP6f events.

To characterize the identity of significantly correlated cells across each behavior state, we first selected cell pairs that are significantly correlated during resting and computed the average correlation of the same cell pairs during running. Similarly, for cell pairs that are significantly correlated during running, we computed the average correlation of these cells during resting. Finally, we calculated the average correlation of the cell pairs that were significantly correlated during both running and resting regardless of the behavior state.

To estimate connectivity among modulated cells, we calculated the number of correlated movement-modulated cell pairs as a fraction of all movement-modulated cell pairs. Similarly, we also calculated the number of correlated non-movement-modulated cell pairs as a fraction of all non-movement-modulated cell pairs.

### Network Analysis

To quantify network connectivity patterns, we calculate closeness centrality similarly to the description in Wuchty and Stadler, 2003^72^. Specifically, for each session, we created an undirected graph using correlated cell pairs during running and an undirected graph using correlated cell pairs during resting. Each cell was considered as a node and each correlated cell pair was connected by an edge. Edge weight is the Pearson correlation coefficient (calculated in the appropriate state, resting or running) between the binarized calcium traces of the cell pair. For each node *i*, closeness centrality *c*(*i*) is defined as:

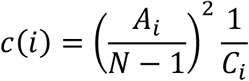

where *A*_*i*_is the number of nodes reachable to node *i* and *C*_*i*_ is the is the sum of distances from node *i* to all reachable nodes. The distance *d*(*i, j*) between nodes *i* and *j* is defined as:

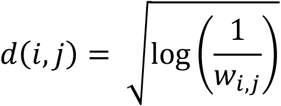

where *w*_*i,j*_ is the edge weight. Closeness centralities of all the nodes were averaged within each network and multiplied by the number of nodes for normalization across networks with different numbers of nodes. Force-directed networks were created using a MATLAB implementation of a force-directed node placement algorithm that spatially clusters nodes proportional to *a*(*i, j*)^73^.

## Acknowledgements

X.H. acknowledges funding from NSF (CBET-1848029, CIF-1955981, DIOS-2002971). R.A.M. acknowledges funding from NIH F31 MH123008-03 and NIH/NIGMS Quantitative Biology and Physiology Fellowship (T32 GM008764) through the Boston University Biomedical Engineering Department. The authors acknowledge computational work done on the Shared Computing Cluster in Boston University’s Research Computing Services. The authors acknowledge research support from the Boston University Micro and Nano Imaging Facility (S10OD024993).

## Author Contributions

Conceptualization, R.A.M. and X.H.; Investigation, R.A.M.; Formal Analysis, M.A.; Resources, M.O. and H.Y.M.; Methodology, A.S.; Software, H.T., S.S., C.Z.; Writing – Original Draft, R.A.M. and X.H.; Writing – Review & Editing, R.A.M., M.A., H.Y.M., X.H.; Supervision – X.H.

