## Supplemental Figure 1 for "The autism spectrum disorder risk gene *NEXMIF* alters hippocampal CA1 cellular and network dynamics"

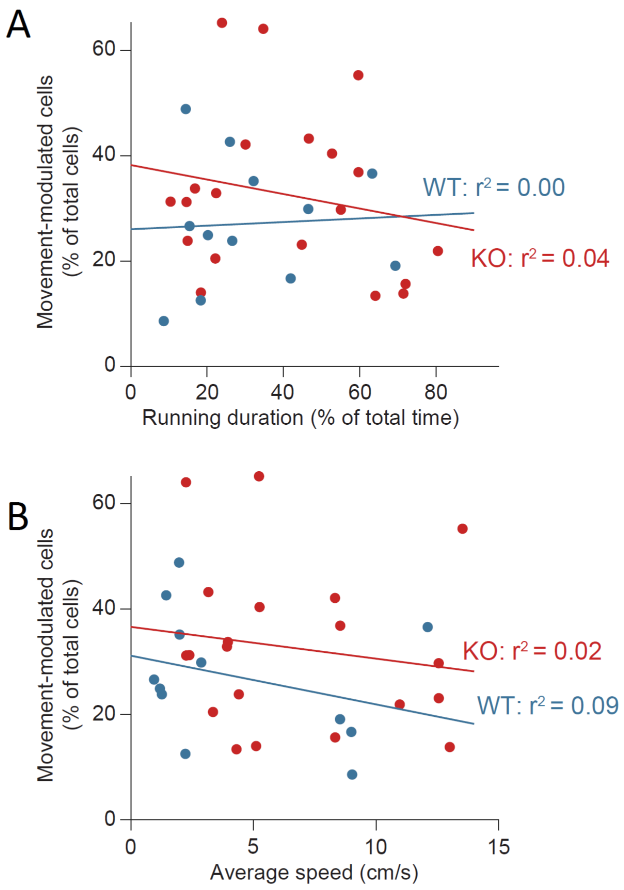


**Figure S1.** Percentage of movement-modulated cells versus (A) movement bout duration and (B) average speed of individual sessions. Linear regression is shown for each population with R-squared value. Bout duration: WT p= 0.87, KO p= 0.40; average speed: WT p= 0.33, KO p= 0.52. n= 12 sessions in 6 WT mice and n= 20 sessions in 8 KO mice.
